# Visuo-motor transformations in the intraparietal sulcus mediate the acquisition of endovascular medical skill

**DOI:** 10.1101/2022.06.15.496236

**Authors:** Katja I. Paul, Karsten Müller, Paul-Noel Rousseau, Annegret Glathe, Niels A. Taatgen, Fokie Cnossen, Peter Lanzer, Arno Villringer, Christopher J. Steele

**Affiliations:** Bernoulli Institute for Mathematics, Computer Science and Artificial Intelligence, University of Groningen, The Netherlands; Mitteldeutsches Herzzentrum, Health Care Center Bitterfeld-Wolfen GmbH, Bitterfeld-Wolfen, Germany; Day Clinic for Cognitive Neurology, University of Leipzig Medical Center, Leipzig, Germany; Department of Neurology, Max-Planck Institute for Human Cognitive and Brain Sciences, Leipzig, Germany; Department of Psychology, Concordia University, Montreal, Canada; Berlin School of Mind and Brain, Humboldt-Universität zu Berlin; Faculty of Medicine, University of Leipzig, Leipzig, Germany; Center for Stroke Research Berlin, Charité Universitätsmedizin, Berlin, Germany

## Abstract

Performing endovascular medical interventions safely and efficiently requires a diverse set of skills that need to be practised in dedicated training sessions. Here, we used multimodal magnetic resonance (MR) imaging to determine the structural and functional plasticity and core skills associated with skill acquisition. A training group learned to perform a simulator-based endovascular procedure, while a control group performed a simplified version of the task; multimodal MR images were acquired before and after training. Using a well-controlled interaction design, we found strong, multimodal evidence for the role of the intraparietal sulcus (IPS) in endovascular skill acquisition that is in line with previous work implicating the structure in simple visuo-motor and mental rotation tasks. Our results provide a unique window into the multimodal nature of rapid structural and functional plasticity of the human brain while learning a multifaceted and complex clinical skill. Further, our results provide a detailed description of the plasticity process associated with endovascular skill acquisition and highlight specific facets of skills that could enhance current medical pedagogy and be useful to explicitly target during clinical resident training.

## Introduction

Since their introduction in 1960 endovascular interventions (EIs) have become one of the principle means to treat cardiovascular diseases ^1^. However, performing these procedures requires a diverse set of skills and extensive practice that is acquired largely through study of literature and observation (knowledge that) and supervised practice (knowledge how). Mastery of EIs requires acquiring generic and task specific skills including visuo-motor skills. The acquisition of these particular skills is demanding due to the minimally invasive nature of the task characterized by manipulating small instruments at long distances via a small incision under the guidance of imperfect imaging systems, typically fluoroscopy (dynamic x-rays). While the endovascular tools are controlled outside the patient, the actual intervention takes place remotely with limited tactile feedback. Since the fluoroscopy x-ray images only provide 2D projections of cardiovascular structures, performing such procedures requires the ability to form a 3D representation of the target site based on multiple projections of 2D images ^2^. Using this mental image as guidance, bi-manual fine motor control and coordination is needed to steer the endovascular tools carefully and effectively through the heart and vascular system. In an earlier publication, we showed that mental rotation ability predicts the learning rate of novices acquiring endovascular skills (Paul et al., 2021). However, insight into the neural correlates of endovascular skill acquisition may provide further insights into the neuro-physiological nature of these skills possibly allowing the development of structured endovascular training curricula.

To date, endovascular skills are acquired by trainees by first observing an expert in the catheter-laboratory followed by gradually performing the procedure under supervision. This means that there is rarely an explicit or structured skills-based training curriculum (Lanzer & Taatgen, 2013). However, operator skills influence the outcome of a procedure and affect patient safety ^4^. In order to develop an explicit training curriculum, insight into the core skills that are required to perform a procedure successfully is needed. Nevertheless, to date little is known about how endovascular skills develop and which factors contribute to safe and efficient performance. Knowledge about which sub-skills are key to the formation of endovascular skills could shape the development of an explicit endovascular training protocol. In this study, we aimed to tease apart the neural correlates and driving force behind learning to perform an EI by examining training-related plasticity in grey matter, white matter microstructure and resting-state functional connectivity.

While research on endovascular skill acquisition with neuroimaging is not yet available, recent work has examined functional magnetic resonance imaging (fMRI) blood oxygen level-dependent (BOLD) signal during and after laparoscopy training, a procedure that is associated with visuo-motor challenges similar to endovascular procedures ^5–8^. Simulator-based laparoscopy training has been found to induce bi-lateral increases in BOLD response in the ventral fronto-parietal grasping network and training-related activity increases in the left M1-hand area predicted participants’ learning rate ^8^. Interestingly, another study showed that the activation pattern while performing simple laparoscopy tasks differed between high -and low-level novice laparoscopy performers. Lower-level performers had a greater BOLD response in the supplementary motor area (SMA) than higher-level performers ^6^. The authors hypothesized that higher activity in the lower-level performers may reflect ongoing, effortful learning, while the necessary motor control might have already become automatic in the higher-level performers. Bahrami and colleagues ^5^ showed that compared to simpler laparoscopic tasks, performing complex laparoscopic tasks led to the recruitment of more brain regions. Simpler laparoscopic tasks activated motor areas (primary motor cortex (M1), supplementary motor area (SMA) and the premotor cortex (PMC)) and the primary somatosensory cortex while the most complex laparoscopic task differentially recruited the superior parietal lobule - possibly reflecting the greater visuo-motor coordination requirements of the more complex task ^5^.

Voxel-based morphometry has been widely used to study changes in grey matter volume (GMV) as a result of short and long-term visuo-motor training, for example, juggling and motor sequence learning ^9^. Structural changes following such training paradigms have been found in the intraparietal sulcus (IPS), mid temporal area (MT/V5), M1, dorsolateral prefrontal cortex (DLPFC) and the ventral PMC ^10–13^. These regions have been associated with various aspects of visuo-motor learning such as the planning and production of movements, motion detection, the integration of motor and sensory information and the storage of their associations ^11,13^. While these tasks share one of the core demands of performing an EI, namely performing visuo-motor transformations, they also differ significantly and are not usually performed within a clinical context.

Diffusion-weighted imaging has also been used to gain insight into training-related changes in white matter microstructure ^14,15^. Often, fractional anisotropy (FA) maps are used as a proxy to identify white matter changes (e.g. Han et al., 2009; Irmen et al., 2020; Jäncke et al., 2009; Scholz et al., 2009; Tremblay et al., 2020). Changes in FA are thought to be reflective of changes in underlying white matter (WM) structure such as myelination, axon density, and thickness ^20^. The only study linking white matter plasticity to skill acquisition in minimally invasive procedures showed that one session of laparoscopy training on a simulator led to a decrease in FA in the superior longitudinal fasciculus adjacent to the ventral PMC region where grey matter changes were detected ^7^. While this study provides solid evidence for WM plasticity over the course of training, the study was not able to identify regions of differential change between the training and control groups that would indicate laparoscopy specific training-related plasticity ^21^.

Resting-state functional connectivity (rs-FC), an MRI technique in which the low frequency fluctuations in BOLD at rest are measured, has been widely used to study functional changes as a result of learning ^22–25^. An advantage of rs-FC over task-based fMRI is that this technique can be used in cases in which it would be difficult to perform the training task inside the scanner e.g., due to space constraints and possible movement artefacts. Rs-FC captures a pattern of co-activation of task relevant networks and, as a result, can be used in a pre/post design to provide insight into underlying functional changes as a result of practising a specific task ^15,23,26^. For example, a study by Albert et al. ^22^ found that 11 minutes of visuo-motor training led to increases in rs-FC in fronto-parietal and cerebellar networks. As these changes were still present after performing an unrelated task, the authors reasoned that the changes in rs-FC may reflect offline processing and memory consolidation. Since then, many studies have provided additional evidence that learning modulates BOLD activity at rest in regions that are associated with the trained task ^7,24,25,27,28^. Hence, rs-FC can shed light on functional networks that are involved in endovascular skill acquisition.

To the best of our knowledge only one study in this research area used an active control group, which was a task-based fMRI study ^8^, and as of yet no study has examined training-related structural plasticity in GMV as a result of a real-life clinical training paradigm. Results of such a study could provide insight into which regions are crucial to perform endovascular procedures and could thereby facilitate the development of targeted training curricula. Furthermore, there is evidence that GMV at baseline can be used to predict the rate of skill acquisition and overall visuo-motor performance ^13,29^. This implies that brain structure before training may be a useful indicator of individual predispositions for successful performance. Not only can GMV change as a result of training, the function of grey matter and white matter microstructure supporting connections between regions can also undergo training-related plastic change. Hence, investigating white matter and functional plasticity should provide additional insights into endovascular skill acquisition.

To examine structural and functional plasticity as a result of learning to perform EIs, we trained medical students naïve to EIs over three days on an endovascular simulator. A separate control group performed the basic first part of this EI to control for visuo-motor execution and the training environment, but controls were not otherwise trained on EI procedures. At baseline, one day prior to training and directly after the last training session, T1-weighted, diffusion-weighted and BOLD EPI scans were acquired from participants in both groups. Based on the current literature, we expected training-related increases in GMV and rs-FC in the PMC, M1 and IPS that were specific to the experimental group and FA changes in close spatial proximity to changes in GMV. Further, we hypothesized that greater GMV in the M1 hand region (M1-hand) at baseline would be predictive of training success. Similar to previous plasticity research on juggling, we also expected that the ability to detect the movement trajectory of the endovascular tools and predict their pathways would be important for performing EIs, so hypothesized that MT/V5 would also exhibit training-related plasticity.

## Method

### Participants

Initially, 42 students in medical studies at the Universities of Leipzig, Halle/Saale and Dresden were recruited from the study programs. Due to technical difficulties with the MRI scanner (not functional on that day) or the simulator (problems with the simulation module), five participants were excluded. The remaining thirty-seven (19 females) participants had a mean age of 23.8 ± 2.55 years. Participants had never performed an EI and had not yet started the practical phase of medical school. We excluded highly skilled musicians, athletes and gamers from the study due their extensive previous visuo-motor practice. All participants were healthy, had normal or corrected to normal vision and had no MRI contraindications. Participants were right-handed as indicated by the Edinburgh Handedness inventory (laterality quotient median 100 ± 10.1, cut-off 64 indicating fully right-handed ^56^). Ethics approval was provided by the ethics committee of the medical faculty of the University of Leipzig, Germany (089/17-ek). All participants signed an informed consent form according to the declaration of Helsinki. At the end of the experiment, participants were reimbursed for their time.

### Experimental procedure

Participants were quasi-randomly assigned to an experimental (9 females, 10 males, 23.9 ±2.54 years) or a control group (10 females, 8 males 23.7 ± 2.64), that were matched as closely as possible for age and gender and trained on an endovascular simulator 80 minutes per day on three consecutive days. The experimental procedure for both groups was exactly the same, only the task performed on the endovascular simulator differed between the two groups. The experimental procedure is shown in Figure 1. In total three MR scans were acquired per participant: at baseline, i.e., two days prior to the pre-scan, one day prior to the training on the simulator (pre-scan) and directly after (within 20 minutes) the last training session on the simulator (post-scan, i.e., 2 days after the pre-scan). Between the pre-scan and the post-scan participants trained on an endovascular simulator.

**Figure 1.**
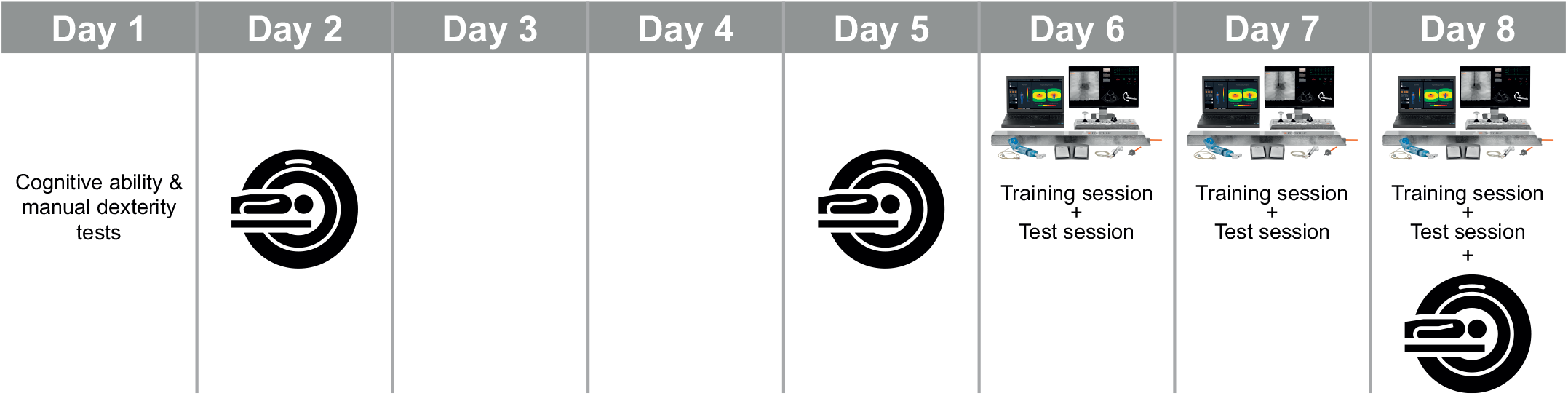
Outline of the experimental procedure. On the first day participants completed cognitive- and a manual dexterity test, on the next day the baseline MR scans (T1, rs-fMRI and DWI) were acquired, after a two day break where nothing study related happened, the MR pre-training scan was performed using the same scanning protocol as for the baseline scan, on the following three days participants either completed the simple training on the endovascular simulator or the complex one followed by a last MR post-training scan after the final simulator session.

### Endovascular simulator

The training took place on the virtual-reality endovascular simulator VIST G5 (Mentice, Gothenburg, Sweden) shown in Figure 2. This simulator included a laptop on which the simulation software was run, a screen on which the simulated x-ray images (fluoroscopy) and simulated vital signs of the patient were depicted, a control panel with a joystick to move the patient table, a foot paddle to control x-ray usage, a syringe to inject contrast agent and a device that depicts the part of the human body in which the tools (catheter and guidewires) were inserted into. Here, the simulated vascular access was at the groin and the tools were inserted via a sheath. The simulator was placed on a table that was adjusted to participants’ height (roughly 15 cm under participants’ elbow). Participants practiced on the simulator while standing. The set-up was designed to simulate how the intervention would be performed in a clinical context.

**Figure 2.**
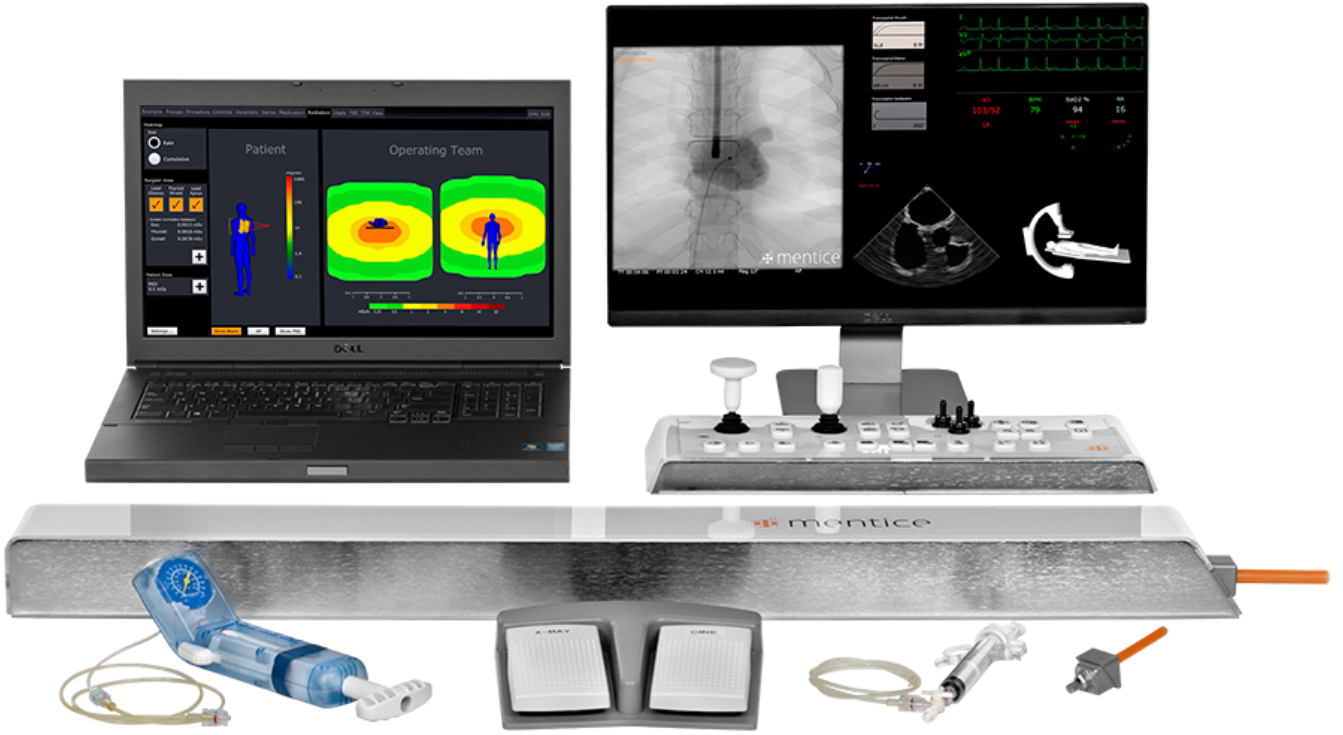
Endovascular simulator VIST G5 (Mentice, Gothenburg, Sweden) including the fluoroscopy screen, human body dummy, syringe to inject contrast agent, control panel to move the patient table and foot paddle to control the x-ray usage.

### Experimental and control tasks

The task of the experimental group on each training day was to perform an angiography of the aortic arch and of the right internal carotid artery. On all three days, participants used a Pigtail catheter to perform the aortic arch angiography. The Pigtail catheter is a standard flush catheter used to inject contrast agent into large arteries such as the aortic arch ^57^. The aortic arch type of the simulated patient varied across days: on day 1, participants trained on an aortic arch type I, on day 2 on a type II arch and on day 3 on a type III arch, respectively. The aortic arch type is defined by the take-off angle of the supra-aortic arteries and determined the type of catheter to be used for cannulating the internal carotid artery. The difficulty to cannulate the artery increases with the aortic arch number. For the type I arch a vertebral catheter was used, for the type II and III arches a Simmons 1 catheter was used. The difference between these catheters is the curvature of their tip designed to facilitate cannulating the target artery. Prior to inserting any of the catheters, a guidewire is inserted over which the catheter is advanced. The guidewire prevents injury of the arteries; moving the catheter within an artery without guidewire support may injure the artery ^57^. In order to visualise the tools within the simulated patient, the participant had to turn on the simulated x-ray via the foot paddle. To keep the instruments in the field of view while advancing the guidewire and catheter to the target position, the participant had to move the patient table via the joystick at the control panel (moving the joystick to the left moved the patient table downwards and brought the lower body part into the field of view, while moving the joystick to the right moved the table upwards). Furthermore, the participant had to turn the C-arm into the LAO (left anterior oblique) 30 position. This position gives an optimal field of view to cannulate the target artery. The angiographies are performed by injecting contrast agent into the target artery followed by the selection of an optimal image. The guidewire and catheters were steered to the target position by pushing /pulling and rotating clockwise or counter clockwise using both hands.

Instead of performing an angiography of the aortic arch and internal carotid artery, the control group only performed the simplified beginning of this procedure. This way, we controlled for the visual-motor executions and the training environment but omitted the complex visuo-motor learning part of the training. Specifically, participants in the control group advanced the guidewire and Pigtail catheter into the proximal part of the aortic arch and then removed the guidewire. On each experimental day, they did this on the respective aortic arch type on which the experimental group trained on that day. The patient table moved automatically and they did not need to rotate the C-arm into the LAO 30-degree position. After conducting this part of the procedure, participants watched a screen capture of the fluoroscopy screen displaying the rest of the procedure that participants in the experimental group actually performed. Rotating the C-arm was not shown on the video to prevent participants from performing this mental rotation step. Participants watched these screen capture videos while standing. This procedure of first conducting the very first part of the procedure on the simulator and then watching the rest of the procedure as a screen capture was repeated on each training day for 80 minutes (i.e., the same total time as the experimental training and test session took). The videos of the shown procedure varied in length and quality.

### Experimental and control group’s training procedure

The experimental group’s training took place individually, and was given by the experimenter who was trained by the manufacturer of the endovascular simulator and by an expert interventional cardiologist with 30 years of experience (PL). On the first training day, participants received written instructions about the task that gave some background information about the task, described its sub-parts, the tools to be used, mentioned clinical guidelines as well as the measured performance metrics. After reading the instructions, an instruction video was shown where an expert (PL) cardiovascular interventional specialist performed the intervention on the endovascular simulator accompanied by audio commentary. Next, the real guidewire and catheter were shown to illustrate the different shapes of the catheter tips and the experimenter explained how to use the simulator. Before starting the first trial, participants were instructed to imagine that the simulator was a real patient, to act as if the simulator was a patient and to take into account that they were novices conducting this procedure for the first time. Specifically, the experimenter highlighted that uncontrolled movements should be avoided and instructed participants to prioritize accuracy over speed.

The experimental groups’ sessions on the simulator always comprised a training and a test session. The training session lasted 60 minutes and the test session 20 minutes. During the training session, verbal feedback was provided that focused on the following aspects: guidewire not guiding the catheter, uncontrolled movements of either of the instruments, incorrect patient table movements, guidewire moving too much while advancing catheter, keeping all instruments under control, spatial orientation of the guidewire and catheter during the procedure, the amount of contrast agent used and the quality of all acquired images. During the test session, participants had 20 minutes to carry out the practised procedure as well as and as often as possible, here no feedback was provided. During the training and test sessions, the fluoroscopy screen was videotaped by the screen capture device Live Gamer Portable 2 ^58^ for later performance evaluations. Furthermore, the simulator automatically captured the total duration of a procedure.

Prior to training on the simulator, control participants also received written instructions and watched an instruction video, both were tailored to the control group’s task. Next, the necessary functionalities of the simulator were explained and the respective guidewire and catheter were shown. While executing the task on the simulator, participants received feedback and the experimenter supervised them while executing the task in the same manner as the experimental group. At the end of the session, participants were asked simple test questions about the watched procedures to increase their motivation to watch the videos and to encourage them to pay attention.

### Performance evaluations

The performance assessment focused on the number of errors committed per procedure and the duration of a procedure. Participants’ performance was retrospectively evaluated using the screen capture videos of the fluoroscopy screen. Two raters first independently counted the number of errors made by participants, and then reviewed and discussed cases where their results were not identical until consensus was reached. The following error types were counted: movement of the catheter ahead of the guidewire; moving the patient table into the wrong direction; accessing the wrong blood vessel with the guidewire and/or catheter; tool not being in the field of view; and inadequate reference picture. The errors committed per simulated procedure were added to yield the total error score per trial. As each day on the simulator comprised a training and a test session that were evaluated separately, there were six simulator sessions. In two out of 114 simulator sessions, a participant did not manage to complete a single trial. Failure to complete a single was penalized by assigning the maximum of the number of errors that was committed by other participants on this trial; for duration the maximum trial time of 20 min was assigned ^37^. The behavioural data for this study has been described in detail in our previous manuscript “Mental rotation ability predicts the acquisition of basic endovascular skills” in which participants’ behavioural performance and the predictability of the learning rate based on cognitive and manual dexterity tests is discussed (Paul et al., 2021).

### Behavioural statistical analysis

We analysed the behavioural data using the statistical software R, version 4.0.0 ^59^. To evaluate whether participants improved over the course of training, linear-mixed effects models were build using the lme4 ^60^ and the lmerTest package ^61^ was used to test the statistical significance of the fixed effects. Two models were built with the dependent variables *mean number of errors* and *mean duration* of a procedure, the fixed effect *session* (i.e., session 1, 2, 3, 4, 5, 6, which refer to the training and respective test sessions) on the simulator and a random intercept for participant (n = 19). As both variables were skewed, they were log-transformed before building the models. We also tested for random slopes to allow for different learning rates across participants. A forward stepwise model fitting procedure was used. As there was only one fixed effect to be modelled, we compared the fixed effect *session* to the null model with only the intercept. We based the model selection for the fixed effect on the p-values. The random effect structure was determined by comparing models using the Likelihood-ratio test; a more complex model was only chosen if it explained significantly more variance. Statistical parameters with a p-value below .05 were regarded as statistically significant.

### MRI acquisition

Data were acquired on a 3T Prisma scanner (Siemens, Erlangen, Germany) with a 32-channel head coil. In all MR sessions, T1-weighed, diffusion weighted and resting-state functional magnetic resonance imaging (rs-fMRI) data were acquired. The scanning protocol for all acquisition days was exactly the same. The T1-weighted structural image was acquired with a MP2RAGE sequence (TR = 5000 ms, TE = 1.96 ms, flip angle = 4°; voxel size = 1 × 1 × 1 mm^3^, ~ 9 minutes duration). Diffusion-weighted data were acquired using a multi-band sequence (66 directions, b = 1000 s/mm^2^, slices = 88, TR = 5200 ms, TE = 75 ms, voxel size = 1.7 mm^2^ isotropic, multi-band acceleration factor = 2, acquisition duration ~ 7 minutes). The rs-fMRI were acquired using a multi-band BOLD EPI-sequence (TR = 1400 ms, TE = 22 ms, flip angle = 67°; voxel size = 2.5 × 2.5 × 2.5 mm^3^, multi-band acceleration factor = 3, ~12 minutes duration). During the rs-fMRI, participants were instructed to look at a fixation cross and to try to not think of anything.

### MRI processing

#### T1-weighted data – Voxel-based morphometry

We used the CAT12 toolbox version 1725 (*CAT12.7 - Computational Anatomy Toolbox for SPM12*, n.d.) to conduct voxel-based morphometry (VBM) which is an extension of the SPM12 ^62^ software running under Matlab R2020b (MathWorks). VBM can measure voxel-wise changes in grey matter concentration ^63^. Data were pre-processed using the standard pre-processing pipeline for longitudinal data to detect small plasticity changes. The steps performed by the pre-processing pipeline included: rigid body registration, bias-correction between time-points, segmentation into tissue classes, shooting spatial registration of the deformed parameters and modulation of the resulting parameters with the Jacobian determinant to compute grey matter volume. The data were smoothed with a Gaussian kernel of 8 mm full width half maximum (FWHM).

#### Rs-fMRI data – intrinsic connectivity

We used the CONN toolbox version 20b running under SPM12 and Matlab R2020b (MathWorks) to calculate intrinsic connectivity (ICC). ICC is a measure of node centrality and is defined as the squared average of the correlation between a given reference voxel and all other voxels ^51^. This approach has the advantage that it does neither require an a-priori defined correlation threshold nor a seed region. As ICC is a network-based summary measure that does not specifically identify functionally connected regions per-se, exploratory follow-up seed-based analyses can be used to explore which networks were altered by the intervention.

Data were pre-processed using the default pre-processing pipeline which included: slice-time correction, realignment and unwarping, motion correction, segmentation into grey matter, white matter and CSF and MNI normalisation. Next, the data were denoised regressing out CSF and WM. The data were filtered with a high-pass filter of 0.01 Hz and smoothed with a Gaussian kernel of 10 mm FWHM.

#### Diffusion-weighted analysis – FA

To analyse the diffusion-weighted data, we used MRtrix ^64^ version 3.0.3 to pre-process and calculate FA maps. Data were first denoised, followed by the recommended DWI general pre-processing pipeline that included correction for current-induced distortion, motion correction and inhomogeneity distortion correction using eddy ^65^ from FSL ^66^ scripted by MRtrix, but dependent on FSL. Next, we estimated the brain mask for each participant and each session, applied bias-field correction and global intensity normalization, fit the diffusion tensor and calculated fractional anisotropy (FA). Co-registration was performed using the advanced normalization tool ^67^ to generate first within-subject’s templates (antsMultivariateTemplateConstruction2.sh: cross-correlation similarity metric and rigid transformation) and then a final template was generated using the output of the within-subjects’ templates as input *antsMultivariateTemplateConstruction2.sh: cross-correlation similarity metric and rigid, 12 dof affine, and SyN nonlinear transformations). This template was then non-linearly registered to the MNI space FA template from FSL to bring it into alignment with the final space maps from CAT12 and CONN. Finally, all transforms were concatenated and applied to the native space FA images to bring them into the common MNI space (antsApplyTransforms). The FA maps were smoothed with a Gaussian kernel of 6 mm FWHM.

### Statistical analysis of MRI data

The focus of the statistical analyses was to determine which, if any, changes could be attributed to the specific effect of training on the endovascular simulator. As such, our main analyses were designed to assess group-specific changes with a GROUP (experimental vs. control group) * TIME (pre- vs. post-training) interaction in each MR modality (VBM, ICC, FA). If a significant interaction effect was identified we followed up with the respective post-hoc tests to specify the presence/direction of change in each group within that modality. Since the rs-fMRI was a network-based metric (ICC), the interaction analysis in this modality was followed up by exploratory seed-based-correlations (SBC) to further specify the functional network of regions involved. Across modalities, we used targeted region of interest (ROI) analyses to identify how regions exhibiting significant changes in one modality may also have undergone plastic change in another.

We also set out to identify the relationship between plasticity and behaviour within the training group. We first correlated mean values from any significant cluster(s) revealed by the interaction analyses with behavioural improvements. Behavioural improvements were calculated as follows: we first calculated accuracy by dividing the number of errors made by each participant per session by the maximum number of errors made by any participant and subtracting that number from 1. The behavioural improvement of each participant was then computed as the increase in accuracy from the first to the last training session. We then followed up with two sets of whole-brain correlation analyses between each metric (VBM, ICC, FA, SBC) and behavioural improvements or overall performance. Overall performance refers to average accuracy across all training sessions. Each of these sets of analyses is described in detail in the following sections. All statistical analyses carried out in SPM were one-tailed, while statistical tests computed in R using extracted mean values were two-tailed tests.

#### Training-specific plasticity – GMV, ICC, FA

Our goal was to investigate whether learning to perform the angiographies led to GMV, ICC and/or FA changes that were specific to endovascular intervention training. To answer this question, we examined the TIME * GROUP interactions with whole-brain GMV, ICC and FA using SPM’s flexible factorial design (n = 37). The interaction analyses were followed up with post-hoc tests to determine the potential nature and direction of any interaction. Any significant clusters identified in the ICC interaction analysis were also followed up by seed-based rs-fMRI analyses to again determine the differential effect of the experimental task over time (TIME * GROUP interaction). All post-hoc analyses were Bonferroni corrected for the four comparisons (i.e, increase or decrease from pre-to post-training in the experimental and the control groups; α = .0125). In all cases, the SPM default threshold on voxel-level p < .001 and cluster-based multiple comparisons correction with FWE (cluster-level p < .05) were regarded as statistically significant. Across-modality ROI analyses were performed using mixed-effect analysis of variance (ANOVA) in R (GROUP * TIME interaction using the package rstatix, Kassambara, 2021).

##### Tissue Mask Generation

In all VBM analyses, a grey matter mask with a threshold of 0.2 was used to prevent bordering effects with white matter or cerebrospinal fluid. In the ICC voxel-wise analysis, we used a mask that was created as follows. The mean of each pre-processed, smoothed rs-fMRI image was computed and a threshold was applied to determine what is inside and outside of the brain. The resulting images were binarized and summed and another threshold was applied as a cut-off to make sure > 99% of images have signal in the resulting area (108 of 109 images). The resulting mask was binarized and multiplied with the standard grey matter mask from FSL to which we applied a threshold of 0.3. Again, the purpose of applying this mask was to prevent partial voluming effects with WM and CSF, while not excluding too many voxels that may actually contain GM. In the FA analysis, we used a group white matter mask generated by calculating the average FA image across participants followed applying a threshold of 0.25 and binarizing the resulting image.

### Relationship Between Plasticity and Behaviour

To investigate whether changes within the experimental group were behaviourally relevant, we correlated the change in mean metric values from any significant interactions with behavioural improvements using Pearson’s correlation (n = 19).

Furthermore, we also used a whole brain approach to test whether individual changes in metrics (GMV, ICC, FA, SBC) of participants in the experimental group were associated with their improvement in performance by correlating the behavioural improvement with voxel-wise change. Finally, we were interested in whether differences in metric values (GMV, ICC, FA, SBC) before training predicted overall performance. Therefore, we tested for a positive correlation between metrics at baseline and the average accuracy per participant across all training sessions. We used this absolute performance measure rather than a rate measure because we were not interested in individual improvement per se, but rather how effective our training intervention was as a whole. In this analysis we corrected for the effects of age and sex and in the VBM analysis for total intracranial volume as well (n = 19).

### Automated meta-analysis of functional correlates of a psychometric predictor of endovascular skill acquisition

In our previous work, we found that mental rotation ability predicted how quickly participants improved across simulator training ^37^. To examine whether networks commonly activated during mental rotation overlap with areas in which we expected to identify structural changes related to endovascular skill acquisition we used Neurosynth ^69^ to conduct an automated meta-analysis of studies that have examined BOLD signal during mental rotation tasks.

### Identification of anatomical regions

Where possible, we used the Anatomy Toolbox version 3.0 ^70^ to identify the location of significant clusters that were revealed by the analyses. The Anatomy Toolbox is an extension of SPM that assigns probabilities that voxels of a given cluster correspond to a certain anatomical region. These maps were created by analysing and mapping the cytoarchitecture of multiple post-mortem brains. However, the atlas does not yet span the entire cortex ^31^.

### Image creation

All images showing brain data were created with the Mango software version 4.1 (Lancaster, Martinez, 2010, http://ric.uthscsa.edu/mango/index.html), figures plotting statistics were created with R version 4.0.0 ^59^.

## Results

### Behavioural results: Learning effects of the experimental group

The two final models consisted of *session* as a fixed effect and a random intercept for participant and either the dependent variable number of errors or the duration of a procedure. The linear mixed effect models indicated a significant learning effect over the course of training. That is, the mean decrease in number of errors (β = -.47, *p* = 2e^-16^) and duration (β = -.21, *p* = 2e^-16^) of a procedure from one session on the simulator to the next was significant. Figure 3 shows a line plot with the number of errors and duration of a procedure per session on the simulator.

**Figure 3.**
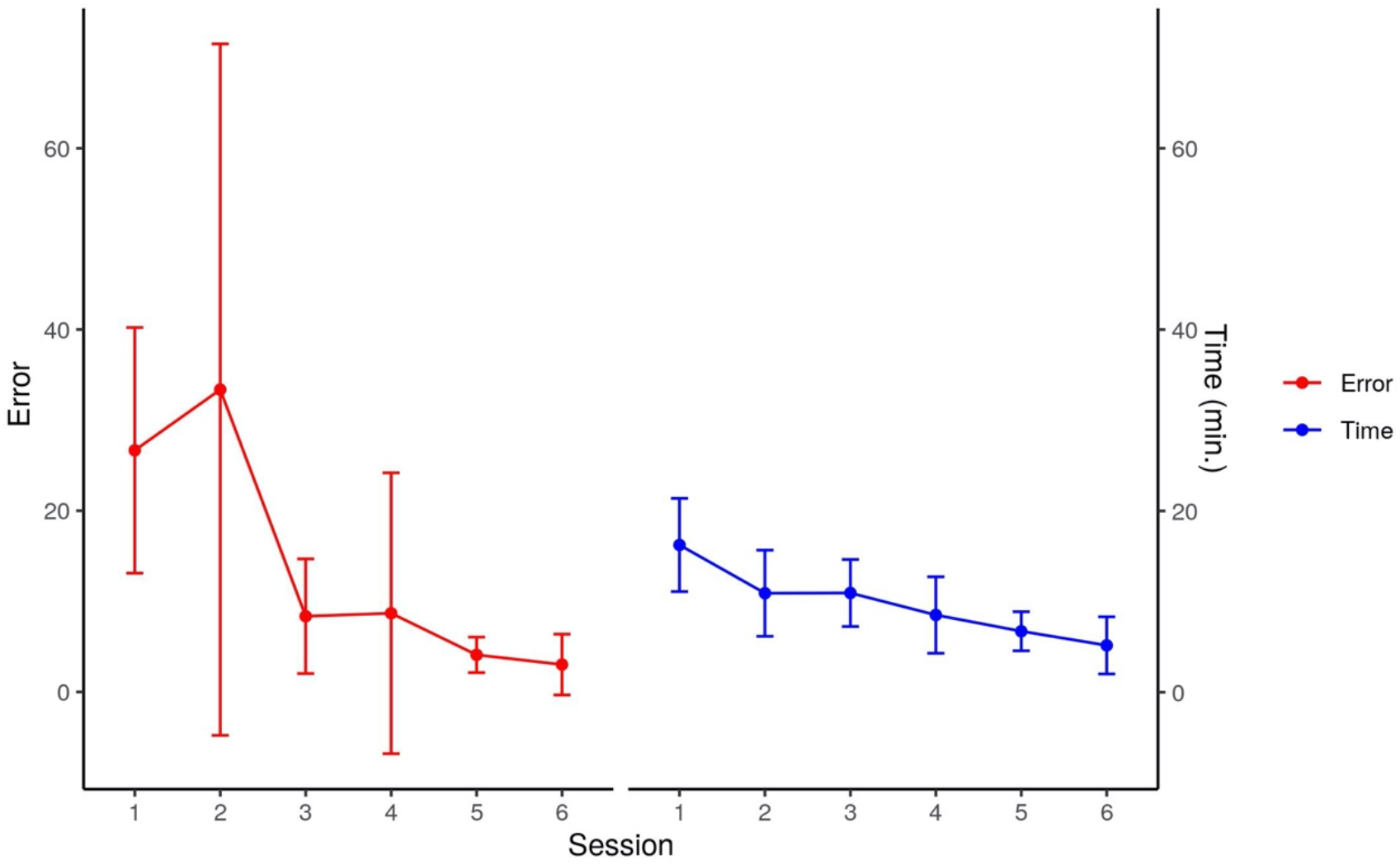
The line plot shows the learning effect of the experimental group across simulator sessions (Training session 1 test session 1, training session 2 test session 2, training session 2 test session 2). The points indicate the mean number of errors and mean duration of a procedure per session and the whiskers indicate the standard deviations.

### Training-specific plasticity: Interaction effect TIME (pre vs. post training) by GROUP (experimental vs. control group)

#### Gray matter volume

The GROUP * TIME interaction revealed two significant clusters in the right hemisphere: in the intraparietal sulcus (IPS, t-value = 4.94, p_FWE-corr_ = .015) and in the primary somatosensory cortex (S1, t-value = 4.51, p_FWE-corr_ = .008), the IPS is depicted in Figure 4a. One cluster in the visual cortex also showed a trend towards statistical significance (t-value = 4.58, *p_FWE-corr_* = .086). Post-hoc tests revealed several significant clusters assessing GMV increase in the experimental group (Table 1). However, only the IPS and visual cortex were also identified in the interaction, indicating that only these clusters showed an increase in GMV that can be specifically attributed to the endovascular training. The identified region of the IPS corresponds to area hlP3, which is the anterior part of the medial wall involved in coordinating reaching movements ^30,31^. The remaining clusters showed an increase in GMV from pre-to post-training in the experimental group, however these increases were not significantly larger than in the control group. The S1 cluster found in the interaction was not confirmed by post-hoc testing and is therefore not discussed any further. None of the remaining post-hoc tests revealed significant effects (p_FWE-corr_ > .05).

**Figure 4.**
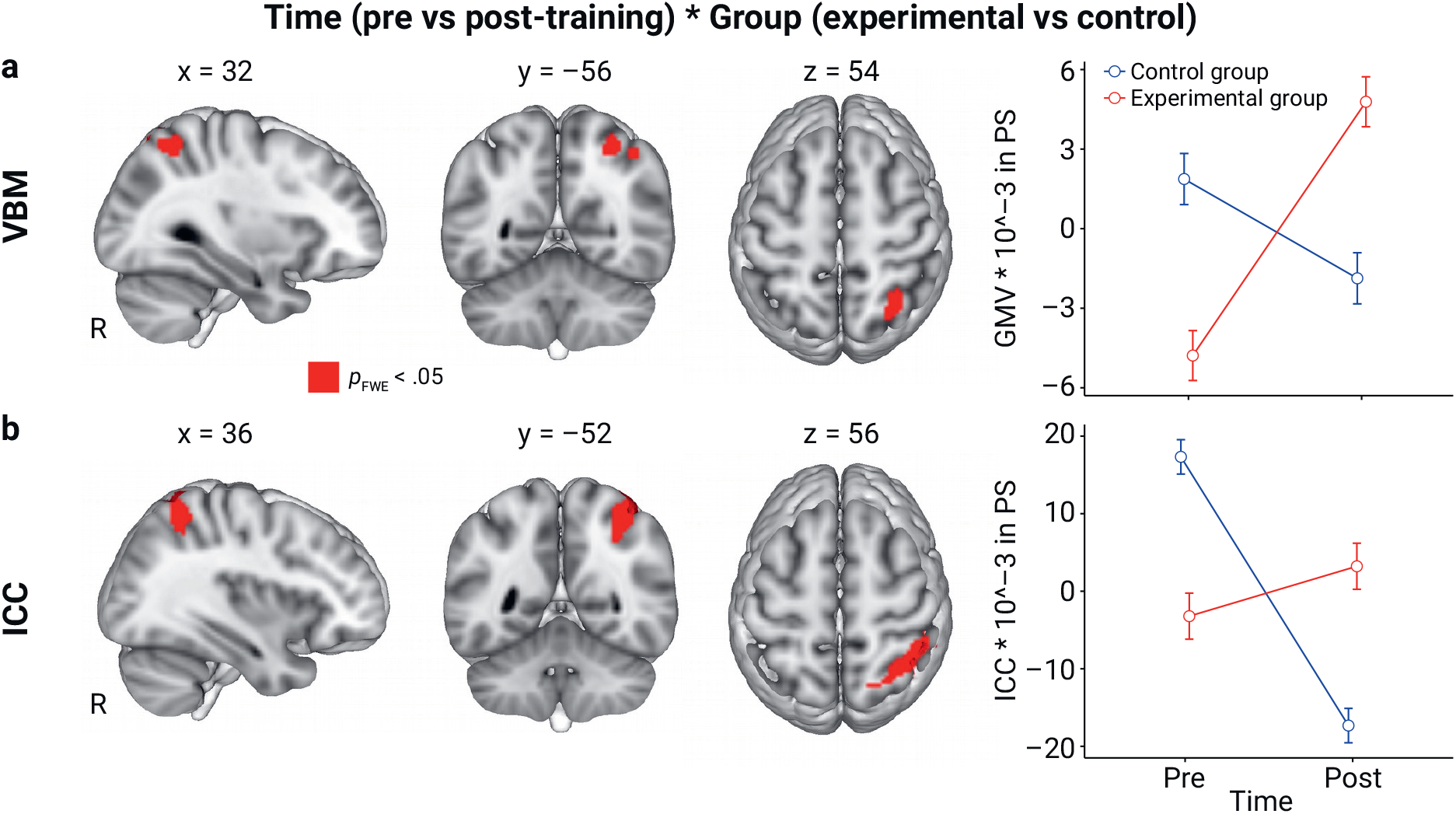
**a)** The figure shows the significant cluster in the right anterior intraparietal sulcus (IPS) revealed by the time (pre- vs post-training) * group (experimental vs control) voxel-based morphometry (VBM) analysis and the interaction plot using centred mean grey matter volume (GMV) values and standard errors **b)** shows the cluster in the right anterior IPS revealed by the intrinsic connectivity (ICC) interaction analysis, the interaction plot shows centred mean ICC values and standard errors. There were no significant baseline differences between groups in neither the VBM nor the ICC analysis (p > .05).

**Table 1.** The table shows the significant clusters revealed by the post-hoc test assessing increases in grey matter volume (GMV) in the experimental group that followed a significant time (pre- vs post-training) * group (experimental vs control) interaction analysis.

| Brain region where change in GMV was found | SPM Anatomy toolbox | Hemisphere | Cluster-level |  |  |  |  |
| --- | --- | --- | --- | --- | --- | --- | --- |
|  |  |  | MNI152 coordinates | Cluster size | t-value | z-value | p-value |
| S1/ BA3b | 3b | r | 50, -18, 38 | 882 | 7.47 | 5.74 | 0.000* |
| Frontal pole | Fo1 | r | 10, 39, -30 | 427 | 6.83 | 5.41 | 0.019 |
| Frontal pole | Fp1 | l | -33, 52, 15 | 752 | 6.56 | 5.26 | 0.001* |
| Inferior lateral occipital Cortex/ V5 | hOc4la | l | -36, -82, 9 | 544 | 6.22 | 5.07 | 0.006* |
| Frontal pole/ Broca's area | BA 45/ OP9 | r | 45, 45, 8 | 1980 | 5.98 | 4.93 | 0.000* |
| Temporal Occipital Fusiform Cortex/V5 | FG4 | r | 50, -56, -6 | 733 | 5.83 | 4.84 | 0.001* |
| Temporal pole | Te3 | r | 54, 9, -4 | 550 | 5.74 | 4.79 | 0.006* |
| Motor cortex/precentral gyrus | 4p | l | -57, 2, 32 | 400 | 5.44 | 4.6 | 0.025 |
| Occipital Fusiform cortex | /hOc1 (V1),<br>hOc2 (V2),<br>hOcV3 (V3v) | l | -10, -64, 3 | 713 | 5.39 | 4.57 | 0.001* |
| Superior parietal cortex/IPS | hIP3/ IPS | r | 32, -58, 52 | 534 | 5.33 | 4.53 | 0.006* |

Note: r indicates right and l indicates left hemisphere. P-values are shown at cluster-level $p_{FWE-corrected} < .05$ . The asterisk \* indicates clusters that are still significant when correcting for conducting four post-hoc tests ( $p_{FWE-corrected} < .0125$ ).

#### Intrinsic connectivity

The GROUP * TIME interaction analysis revealed a significant cluster in the right IPS (t-value = 5.54, p_FWE-corr_ = .006, shown in Figure 4b) that largely overlapped with the significant cluster from the VBM interaction analysis (Figure 4a). A post-hoc test revealed that ICC in the right IPS in the control group was larger before training compared to after training. Three other regions were revealed by the same post-hoc test listed in Supplementary Table S1; however, we cannot attribute these changes specifically to the control task because they were not present in the interaction. None of the remaining post-hoc tests revealed any significant changes in ICC over time (*p_FWE-corr_* > .05).

#### Fractional anisotropy

The whole-brain GROUP * TIME interaction analysis did not reveal any significant effects (*p_FWE-corr_* > .05).

##### ROI-analysis of FA surrounding the cluster revealed by the VBM interaction analysis (IPS)

A ROI analysis using the cluster in the IPS revealed by the VBM analysis revealed a significant TIME * GROUP interaction (*F* (1, 35) = 4.43, *p* = .043, Figure 5). The post-hoc tests indicated that the interaction was driven by a decrease in the control group (experimental group: *t* = −1.08, *p* = .29; control group: *t* = 2.14, *p* = .047).

**Figure 5.**
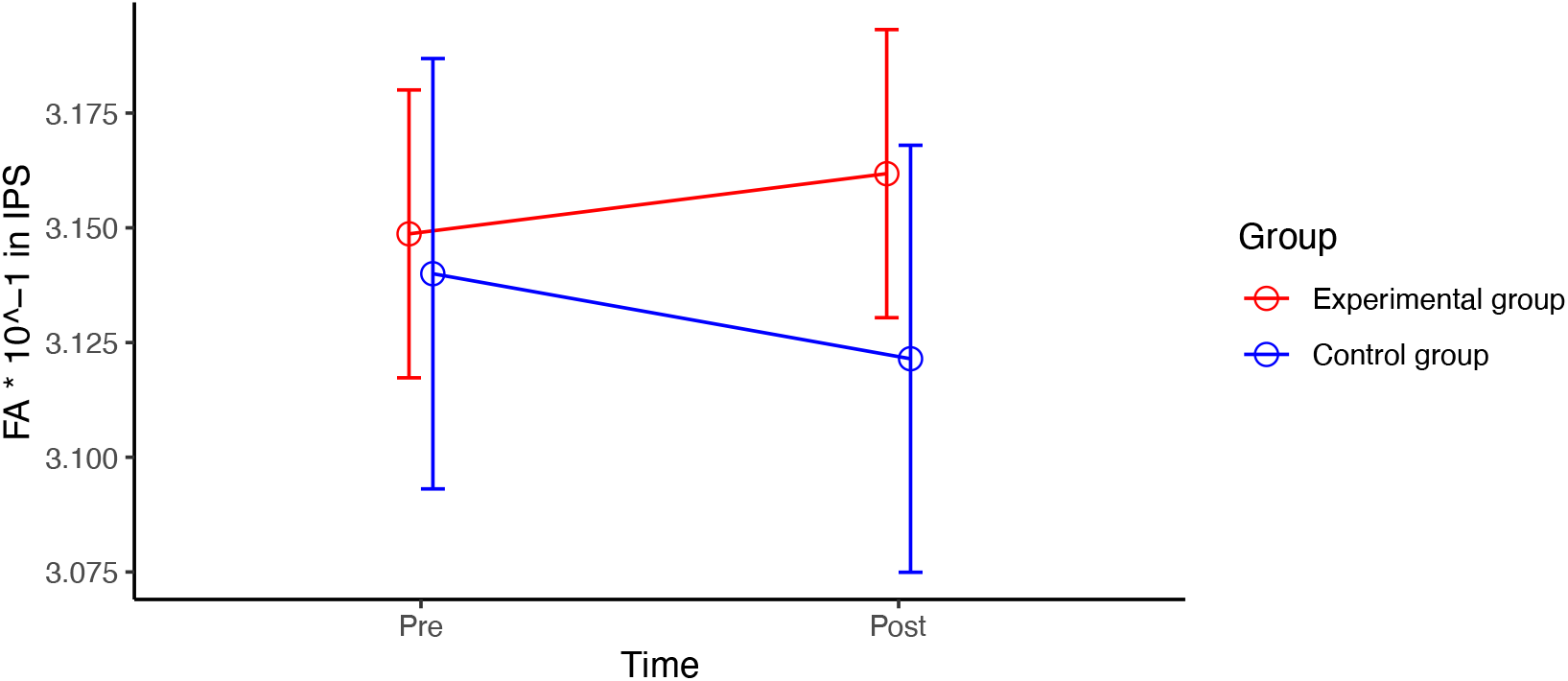
Interaction plot showing mean fractional anisotropy (FA) values extracted from the region of interest in and around the white matter of the IPS cluster revealed by the VBM time (pre- vs post-training) * group (experimental vs control) interaction analysis.

##### Targeted correlations with behaviour

To test whether training-related changes identified by the GMV, ICC, and FA interaction analyses were behaviourally relevant, we correlated the changes in the parameters within the identified significant clusters with behavioural improvement. None of the correlations were statistically significant (*p* > .05).

#### Seed-based correlation: IPS

The ICC analysis identified group-specific changes in rs-FC in the IPS. To identify how functional connectivity to the IPS changed over the course of training we conducted an exploratory SBC with the IPS as the seed. The GROUP * TIME interaction analysis revealed the following seven clusters: orbitofrontal cortex, Crus I, bilateral IPS, frontal pole, precentral gyrus and inferior temporal gyrus (Figure 6, Supplementary Table S2). Post-hoc testing showed that the cluster in the left IPS and the precentral gyrus were driven by increases from pre- to post-training in the experimental group (Supplementary Table S3). These two regions are associated with motor planning and visuo-motor integration and may suggest ongoing learning in the experimental group who practised the complex task on the simulator ^6,31–33^. The cluster in the left Crus I of the cerebellum was confirmed by the contrast control group pre > post-training (t-value = 6.47, p_FWE-corr_ = .001). This means that connectivity between the IPS and the left Crus I decreased over time in the control group. Since Crus I has been shown to be involved in higher cognitive functions, such as working memory and attention, we interpret this decrease in connectivity between the IPS and Crus I as fewer attentional resources being needed for the visuo-motor integration after training in the control group ^34^.

**Figure 6.**
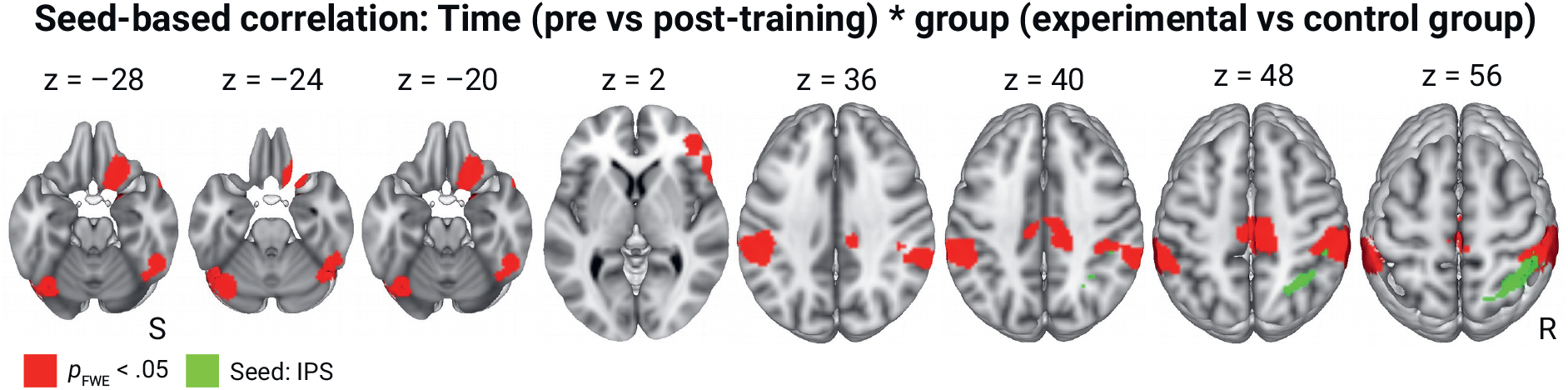
The axial slices show the clusters revealed by the time (pre- vs post-training) * group (experimental vs control) interaction seed-based correlation analysis with the IPS as the seed received from the significant ICC time (pre- vs post-training) * group (experimental vs control) interaction analysis. More detailed information on these clusters can be found in Supplementary Table 2.

### Relationship Between Plasticity and Behaviour

The following sections list the results from the whole-brain correlational analyses that tested for an association between changes from pre- to post-training in GMV, ICC, FA and SBC and individual behavioural performance improvements in the experimental group.

#### Gray matter volume

Correlating the behavioural improvement with the change in GMV using the whole brain approach revealed a significant positive correlation in a cluster spanning V1V2V3 in the left and right hemisphere (V1V2V3, t-value = 6.17, p_FWE-corr_ = .007, see Figure 7). Interestingly, this cluster largely overlaps with the one that showed a trend towards statistical significance in the VBM interaction analysis and was found to be significant in the post-hoc test (see Figure 7c). This result indicates that changes in the brain regions that are associated with visual information processing scaled with behavioural improvements. There were no significant negative correlations.

**Figure 7.**
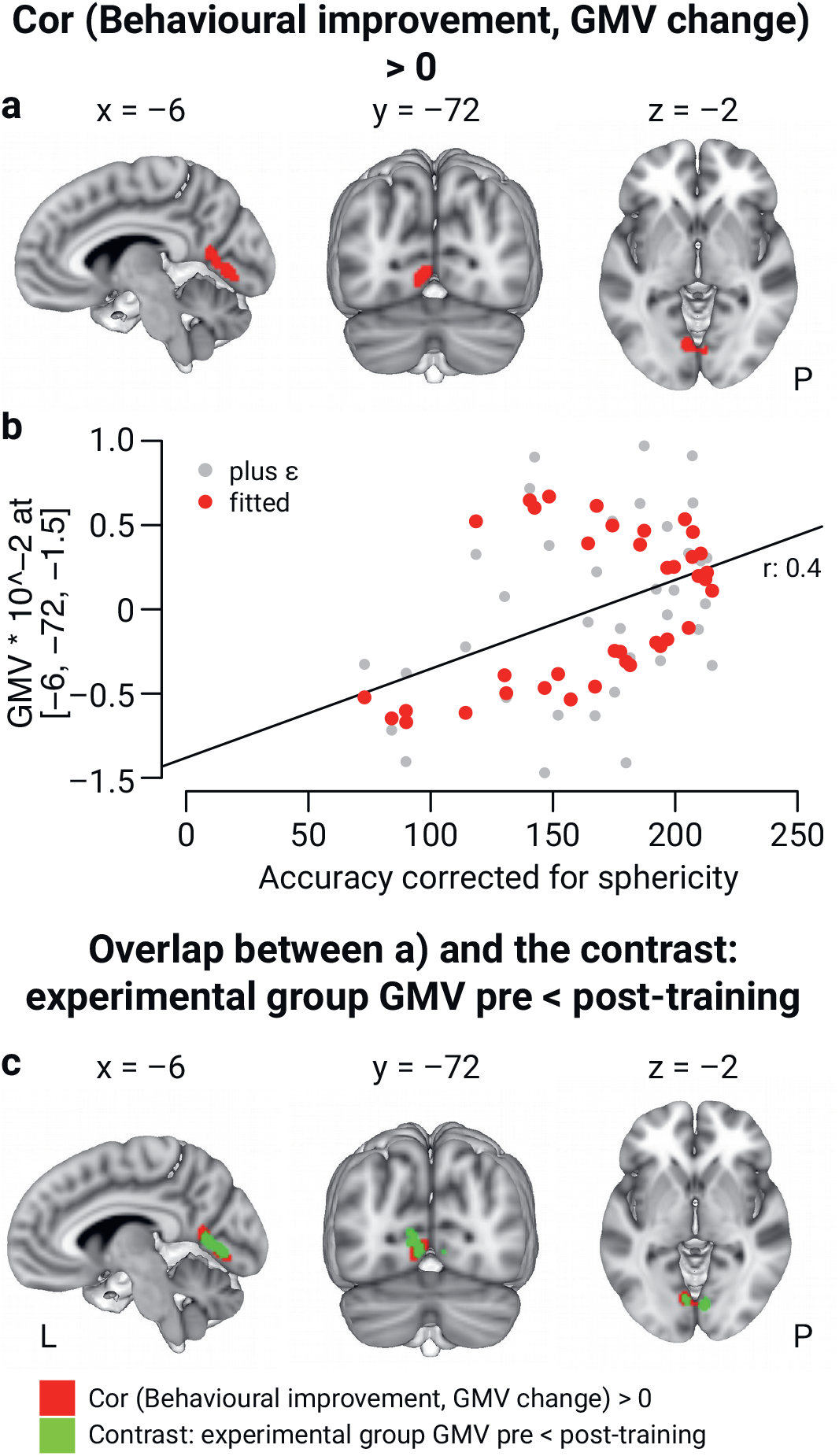
**a)** The figure shows the cluster in visual areas V1V2V3 spanning the right and left hemisphere revealed by correlating behavioural improvements with GMV change in the experimental group. **b)** The scatterplot shows the correlation between behavioural improvements and average GMV changes from pre-to-post-training in V1V2V3. The values are sphericity corrected. Fitted refers to GMV values within the statistical model, while plus epsilon refers to centred GMV values. **c)** This part of the figure shows that the significant cluster revealed by the correlational analysis shown in Figure 7a) largely overlaps with the cluster revealed by the post-hoc contrast assessing GMV increases in the experimental group. The correlation coefficient r indicates the strength of the correlation using the plus epsilon values and is only shown for visualisation purposes and not meant for interpretation.

#### Intrinsic connectivity and fractional anisotropy

Neither changes from pre- to post-training in ICC nor FA in the experimental group correlated with the behavioural improvement (p_FWE-corr_ >.05).

#### Seed-based correlation: IPS

We could not identify any significant correlations between changes in SBC between the IPS and any other brain regions in the experimental group and behavioural improvements (p_FWE-corr_ >. 05).

#### Seed-based correlation: V1V2V3

Exploratory correlational analysis with the SBC in the experimental group between V1V2V3 and the functional activation of the rest of the brain with the behavioural performance improvement identified a positive correlation bilaterally in the middle temporal gyrus (MTG) and bi-laterally in the IPS (see Figure 8). That is, larger improvements across training were associated with an increase in rs-FC in these regions. In particular, the increase in connectivity between the IPS and V1V2V3 is interesting because the same region exhibited changes in GMV and rs-FC in the GROUP * TIME interaction analyses. These results indicate that behavioural improvement was associated with increased correlation between V1V2V3 and bilateral IPS and MTG at rest. The IPS receives input from V1V2V3 for visuo-motor coordination; the MTG is associated with multimodal sensory integration ^35^. The analysis did not reveal any negative correlations.

**Figure 8.**
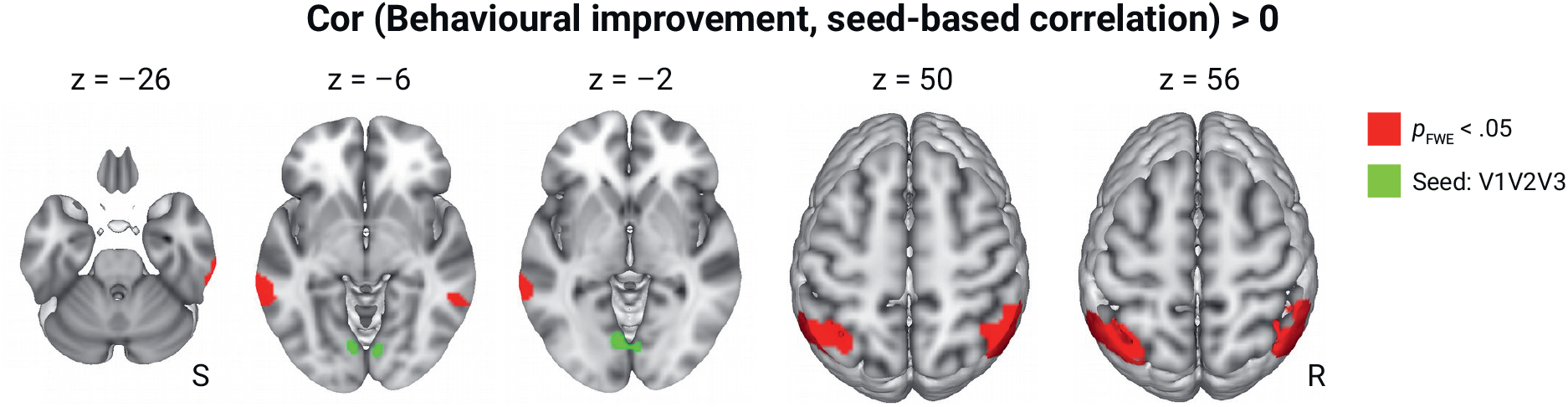
Seed-based correlation between a cluster in the visual cortex (V1V2V3) and functional activity in the rest of the brain. The cluster V1V2V3 was derived by correlating the behavioural improvement with grey matter volume increases in the experimental group. The axial slices show the clusters revealed by a positive correlation, i.e., a larger increase in accuracy is associated with a larger increase in connectivity between V1V2V3 and the shown clusters in the bi-lateral middle temporal gyrus and bi-lateral IPS.

### Predicting performance from baseline GMV, ICC, FA and SBC

In the following analyses, we tested whether individual differences in GMV, ICC, FA and SBC at the baseline scan prior to training could predict participants’ overall performance across simulator training.

#### Gray matter volume

GMV at baseline in lobule VIIIb of the cerebellum (t-value = 5.22, p_FWE-corr_ = .034) in the right hemisphere positively correlated with the average accuracy of participants during the simulator training (see Figure 9). Lobule VIIIb is associated with finger and hand movements as well as tactile representations ^34,36^. There were no significant negative correlations (p_FWE-corr_ > .05).

**Figure 9.**
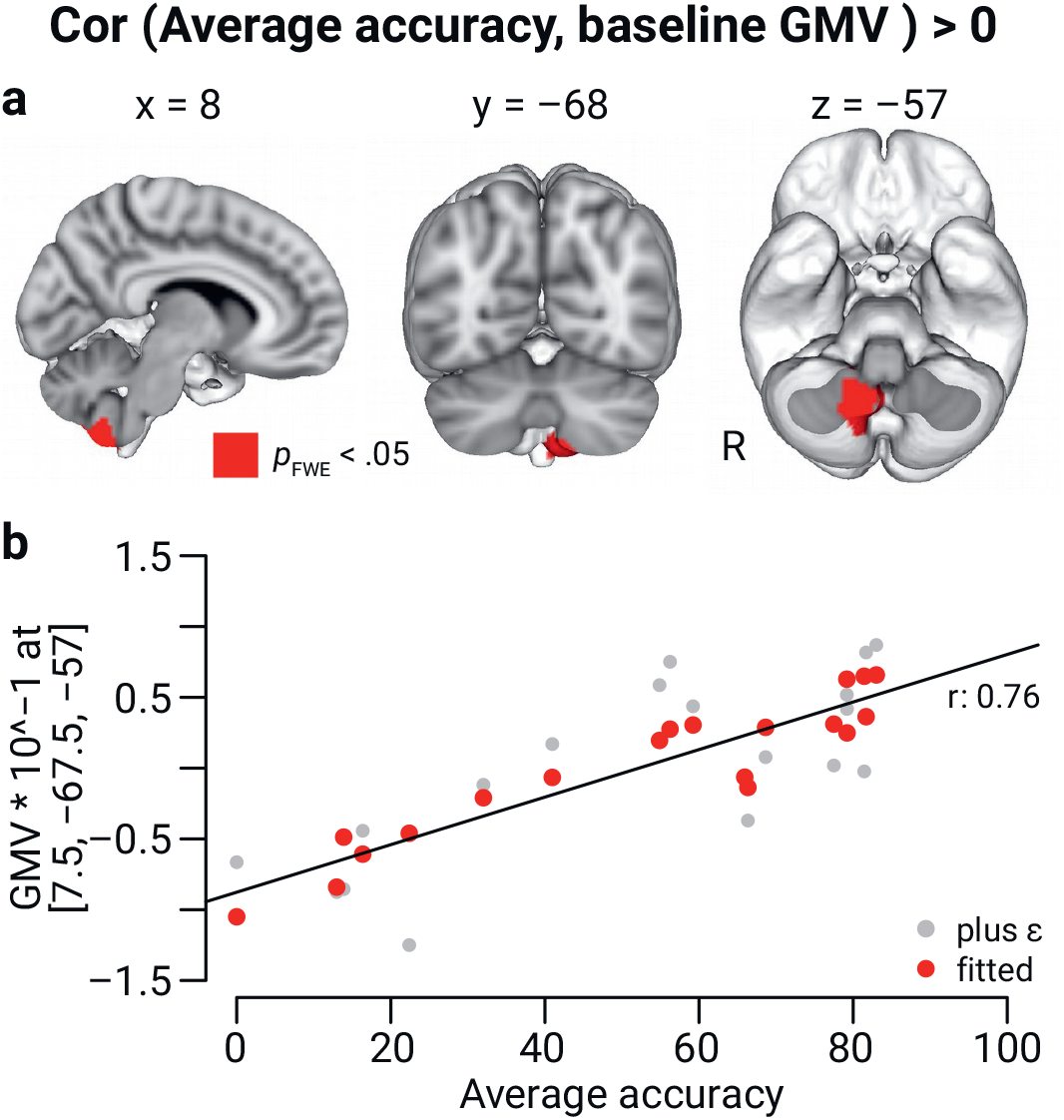
**a)** This figure shows the cluster in Lobule VIIIb in the right cerebellum (MNI 152 coordinates: 8, −68, −57) revealed by correlating GMV at baseline (before training) with overall simulator performance **b)** The scatterplot shows the correlation between average GMV in Lobule VIIIb at baseline and average accuracy across simulator training. Fitted refers to GMV values within the statistical model, while plus epsilon refers to centred GMV values. The correlation coefficient r indicates the strength of the correlation using the plus epsilon values and is only shown for visualisation purposes and not meant for interpretation.

#### Intrinsic connectivity, fractional anisotropy and seed-based correlations

Neither ICC, FA nor any seed-based correlation to the IPS, Lobule VIIIb or V1V2V3 at baseline could predict overall performance on the endovascular simulator (p_FWE-corr_ > .05).

### Automated meta-analysis of functional correlates of a psychometric predictor of endovascular skill acquisition

The automated-meta-analysis of networks activated during mental rotation, a psychometric predictor of EI skill acquisition ^37^, showed that these networks include the right IPS and overlap with the cluster identified in the GROUP * TIME interaction analyses.

## Discussion

The goal of the current paper was to determine which structural and functional plastic changes drive endovascular skill acquisition. Using voxel-based morphometry (VBM), diffusion tensor imaging (DTI) and resting-state fMRI (rs-fMRI), we found multimodal evidence for the involvement of the IPS in endovascular skill acquisition which is consistent with previous plasticity work linking the IPS to visuo-motor skill acquisition including visuo-motor transformations and coordination ^8,11,18,30,38,39^. Our finding extends the literature by being the first strongly controlled, multimodal evidence for the crucial role of this brain structure in a real-life visuo-motor clinical training paradigm. Changes in GMV in multiple areas of the visual cortex, but not the IPS, were associated with participants’ behavioural improvement. However, our exploratory seed-based correlations (SBC) with behavioural improvement identified significant associations between the visual cortex and a bi-lateral clusters in the IPS, supporting our main findings that the IPS is crucial for endovascular skill acquisition. Finally, we also found that individual differences in GMV in Lobule VIIIb before learning predicted participants’ overall performance. These results shed light on the plasticity mechanisms underlying endovascular skill acquisition and may have important implications for training residents in interventional procedures.

### Training-related structural and functional plasticity in the IPS supporting visuo-motor skill acquisition

Our results provide strong evidence of the role of the IPS in basic endovascular skill acquisition. Further supporting our results, a task-based laparoscopy training study by Karabanov and colleagues ^8^ also found functional changes in the IPS with fMRI. In addition, the change we identified in the IPS is backed up by other modalities (ICC and FA), thus providing a complete overview of the plasticity process associated with endovascular skill acquisition. More specifically, we identified changes in the anterior/medial portion of the IPS which transforms visual information for use by the motor regions and thus promotes hand-eye coordination and planning of movements ^30,32^. This function of the IPS fits well with what participants in the experimental group needed to learn, i.e., to read the fluoroscopy images and use this information to coordinate the fine-grained motor movements to steer the endovascular tools through the vascular system. Further supporting this interpretation, we also found independent meta-analytic evidence that the IPS is related to mental rotation ability – which we previously identified as a predictor of behavioural improvements in the same participant sample ^37^. Mental rotation ability is a necessary component of this task, as the fluoroscopy images lack spatial cues. Therefore, the operator first needs to create a mental 3D model of the arteries and mentally manipulate this model to infer the orientation of the tools and curvature of the arteries to steer the tools safely through the blood vessels ^37^. Together with the multimodal evidence for the role of the IPS in the current study, these findings highlight that visuo-motor coordination and transformation (including mental rotation) are core skills for learning and performing endovascular interventions that are supported by the structure and function of the IPS.

### Time-course of training-related structural and functional changes

We found the same pattern of increases/decreases and between-group interactions in all of our MRI metrics (see Figure 4 and 5). As the task of the control group was included in the experimental group’s task, and this group performed more complex steps on top of the control task, we conclude that the net effect of training-related plasticity was an increase in GMV, ICC and FA in the IPS. In contrast, practising only the control task led to decreases in the respective metrics. This pattern may be explained by the expansion-renormalisation model of neural plasticity ^40^. According to this model, during the early stage of learning many changes occur e.g., on a cellular and functional level, later the most suitable change is selected in terms of neural efficiency while the others are eliminated. Applied to our findings, this suggests that participants in the experimental group who learned the complex procedure on the simulator are still in the expansion phase, while the participants in the control group have already acquired the simple task and are thus in the elimination phase. Furthermore, our findings extend the literature by providing evidence for rapid, specific structural brain changes due to only three days of skills training. So far, such rapid remodelling has mostly only been shown in the animal literature ^41,42^.

### Regions contributing to visuo-motor coordination

Increases in GMV in early visual cortical areas (V1V2V3) correlated with behavioural performance improvements. These regions are involved in basic visual information processing, which is a prerequisite for successful visuo-motor coordination in the endovascular task ^43,44^. It is interesting that plasticity in visual cortical regions was associated with learning rather than in the IPS. However, regions V1V2V3 are part of the dorsal stream and project to the IPS and thus are connected to the area where we identified training-related structural and functional changes. The dorsal stream is associated with visually-guided grasping movements and thus essential to performing the movements required by the experimental task ^35,45^. Intriguingly, we also found that increases in rs-FC between V1V2V3 and the bi-lateral IPS were associated with performance changes over the simulator training course. Thus, this result confirms on a functional level that increased coupling between these regions of the dorsal stream is important to endovascular skill acquisition.

### Individual differences in brain structure and function and their relation to endovascular skill acquisition

GMV in right Lobule VIIIb of the cerebellum before training predicted how well participants performed across simulator training. Lobule VIIIb has been shown to be activated during right-handed finger/hand movements and tactile stimulation, and is functionally connected to the superior parietal lobule and sensori-motor regions ^34,36,46^. While visuo-motor transformations as indicated by the change in GMV, FA and ICC in the IPS were related to learning, our findings may indicate that cerebellar somatosensory representations are crucial to overall training success, which is consistent with its role in error correction ^36,47^. This may possibly indicate the importance of a tactile representation of the guidewire to be able to control its movements successfully. The dominant-in our participants right hand-is typically used to control the guidewire while the left hand supports the endovascular tools ^48^. Inter-individual differences in GMV in Lobule VIIIb before training might be caused by differences in previous motor experience or genetic predispositions ^13^. Thus, this finding shows that interindividual differences in GMV are predictive of successful early endovascular skill acquisition.

### Mechanisms underlying GMV, rs-FC and FA changes

Though it is difficult to tie non-invasive MRI measurements of plasticity to underlying physiological mechanisms, invasive histology in non-human animals provides a useful comparison for our findings ^49^. Increases in GMV have been tied to expanding underlying cellular structures such as dendritic spine remodelling in response to new task demands ^20,29,42^. Changes in dendritic spine density have been found after short training periods and may be reflected in the intensity changes in GMV that we detected using VBM ^20,42^. Increased FA indicates an increase in water restriction in white matter, possibly caused by increases in axon density or myelination which could have contributed to learning by facilitating signal transmission ^20,50^. Changes in ICC reflect changes in the correlation of low-frequency BOLD fluctuations between different brain regions. This means that the strength of the connection of an identified region to other voxels in the brain has changed ^51^.

In accordance with similar studies (e.g. Irmen et al., 2020; Karabanov et al., 2019; Taubert et al., 2010) our data show that structural and functional changes associated with skill acquisition occur in the same temporal and spatial domain. Other work ^18^ however found that structural and functional changes don’t happen in parallel. This divergence may be due to differences in training time and paradigm.

### Implications for interventional specialties

Our findings may have important implications for training residents in interventional procedures. They highlight the importance of visuo-motor coordination and mental rotation ability in endovascular skill acquisition and, importantly for medical pedagogy, they provide evidence that only three days of dedicated simulator training guided by focused instruction and specific feedback can lead to performance improvements and structural and functional brain changes. Based on our findings, we suggest that before training on patients, residents complete dedicated, explicit simulator training focused on mental rotation ability and visuo-motor coordination. For example, explicitly practising how to interpret the fluoroscopy images and coordinate endovascular tool manipulation based on this information may facilitate acquiring these basic, yet critically important skills that are a prerequisite to more advanced skills such a decision-making under uncertainty and devising treatment plans. Furthermore, the finding that baseline GMV can predict overall simulator performance suggests that improvements may be affected by pre-existing abilities. Indeed, even at this basic level of EI skill acquisition marked inter-individual differences were observed in novice performers ^37^. Though this observation is interesting from a neuroscience perspective, its relevance in the context of medical education is unclear. Before generating explicit training recommendations based on our experimental findings, we must also understand how aptitude and deliberate practice interact to influence the learning of specific EI skills. While aptitude testing could be used to tailor training to individual needs, at the current state it is impossible and it thus would be unethical to select trainees based on anatomical or functional MR data.

### Limitations

One limitation of our study is that we evaluated performance based on the number of errors participants made during the training, we did not take time taken to complete a procedure into account. This choice was made based on the hypothesis that the number of errors most accurately reflects participants’ visuo-motor performance. The time taken to complete the procedure can be a misleading measure of performance as faster is not always better in the current task, and participants were instructed to prioritise accuracy over speed during the training.

Another potential limitation is that the simulation environment may not replicate all EI skills required in in-vivo settings. However, simulator training provides the best available option to train and to test EI skills without any risk to harm patients. Moreover, the face, content and construct validity of the used simulator has been proven and learning possibly transfers to the clinic ^53–55^.

### Conclusions

In the present study, we have found evidence that parieto-occipital regions are crucial to endovascular skill acquisition and performance. To the best of our knowledge, this is the first study providing multimodal, controlled evidence for plastic changes in the IPS as a result of a real-life visuo-motor clinical training paradigm. Further, we found that structural changes associated with skill acquisition are paralleled by functional ones in the temporal and spatial domain and adhere to predictions by the expansion-renormalisation model. Baseline grey matter volume predicted participants’ absolute performance pointing to individual difference factors that also influence endovascular skill acquisition. Together, these findings shed light on the dynamics of structural and functional plasticity resulting from visuo-motor learning. Moreover, the results point to specific skills that trainees in interventional cardiovascular medicine could practice on a simulator to be later transferred to clinical practice.

## Author contributions

**KIP** drafted the manuscript

**KIP, PL, CJS, FC and AV** were involved in the conception and design of the work

**KIP and AG** acquired the data

**KIP, AG, NAT and PNR** analysed the data

All authors interpreted the data and revised the manuscript

All authors have read and approved the manuscript

## Data availability statement

The datasets generated during and analysed during the current study are available from the corresponding author on reasonable request.

## References

1. Faxon, D. P. & Williams, D. O. Interventional Cardiology: Current Status and Future Directions in Coronary Disease and Valvular Heart Disease. Circulation 133, 2697–2711 (2016).

2. Lanzer, P. Cognitive and Decision-Making Skills in Catheter-Based Cardiovascular Interventions. in Catheter-Based Cardiovascular Interventions: A Knowledge-Based Approach (ed. Lanzer, P.) 113–155 (Springer, 2013). doi:10.1007/978-3-642-27676-7_10.

3. Niels Taatgen, P. L. Procedural Knowledge in Percutaneous Coronary Interventions. J. Clin. Exp. Cardiol. (2013) doi:10.4172/2155-9880.S6-005.

4. Lin, P. H. et al. Carotid artery stenting with neuroprotection: assessing the learning curve and treatment outcome. Am. J. Surg. 190, 855–863 (2005).

5. Bahrami, P. et al. Neuroanatomical Correlates of Laparoscopic Surgery Training. Surg. Endosc. 28, (2014).

6. Garbens, A. et al. Brain activation during laparoscopic tasks in high- and low-performing medical students: a pilot fMRI study. Surg. Endosc. 34, 4837–4845 (2020).

7. Irmen, F. et al. Functional and Structural Plasticity Co-express in a Left Premotor Region During Early Bimanual Skill Learning. Front. Hum. Neurosci. 14, 310 (2020).

8. Karabanov, A. N. et al. Getting to grips with endoscopy - Learning endoscopic surgical skills induces bi-hemispheric plasticity of the grasping network. NeuroImage 189, 32–44 (2019).

9. May, A. Experience-dependent structural plasticity in the adult human brain. Trends Cogn. Sci. 15, 475–482 (2011).

10. Boyke, J., Driemeyer, J., Gaser, C., Buchel, C. & May, A. Training-Induced Brain Structure Changes in the Elderly. J. Neurosci. 28, 7031–7035 (2008).

11. Draganski, B. et al. Changes in grey matter induced by training. Nature 427, 311–312 (2004).

12. Driemeyer, J., Boyke, J., Gaser, C., Büchel, C. & May, A. Changes in Gray Matter Induced by Learning—Revisited. PLOS ONE 3, e2669 (2008).

13. Gryga, M. et al. Bidirectional gray matter changes after complex motor skill learning. Front. Syst. Neurosci. 6, 37 (2012).

14. Chang, Y. Reorganization and plastic changes of the human brain associated with skill learning and expertise. Front. Hum. Neurosci. 8, 35 (2014).

15. Taubert, M., Villringer, A. & Ragert, P. Learning-Related Gray and White Matter Changes in Humans: An Update. The Neuroscientist 18, 320–325 (2012).

16. Han, Y. et al. Gray matter density and white matter integrity in pianists’ brain: A combined structural and diffusion tensor MRI study. Neurosci. Lett. 459, 3–6 (2009).

17. Jäncke, L., Koeneke, S., Hoppe, A., Rominger, C. & Hänggi, J. The Architecture of the Golfer’s Brain. PLoS ONE 4, e4785 (2009).

18. Scholz, J., Klein, M. C., Behrens, T. E. J. & Johansen-Berg, H. Training induces changes in white-matter architecture. Nat. Neurosci. 12, 1370–1371 (2009).

19. Tremblay, S. A. et al. White matter microstructural changes in short-term learning of a continuous visuomotor sequence. http://biorxiv.org/lookup/doi/10.1101/2020.10.02.324004 (2020) doi:10.1101/2020.10.02.324004.

20. Zatorre, R. J., Fields, R. D. & Johansen-Berg, H. Plasticity in gray and white: neuroimaging changes in brain structure during learning. Nat. Neurosci. 15, 528–536 (2012).

21. Thomas, C. & Baker, C. I. Teaching an adult brain new tricks: A critical review of evidence for training-dependent structural plasticity in humans. NeuroImage 73, 225–236 (2013).

22. Albert, N. B., Robertson, E. M. & Miall, R. C. The Resting Human Brain and Motor Learning. Curr. Biol. 19, 1023–1027 (2009).

23. Biswal, B., Yetkin, F. Z., Haughton, V. M. & Hyde, J. S. Functional connectivity in the motor cortex of resting human brain using echo-planar mri. Magn. Reson. Med. 34, 537–541 (1995).

24. Gong, D. et al. Enhanced functional connectivity and increased gray matter volume of insula related to action video game playing. Sci. Rep. 5, 9763 (2015).

25. Sampaio-Baptista, C. et al. Changes in functional connectivity and GABA levels with long-term motor learning. NeuroImage 106, 15–20 (2015).

26. Guerra-Carrillo, B., Mackey, A. P. & Bunge, S. A. Resting-State fMRI: A Window into Human Brain Plasticity. The Neuroscientist 20, 522–533 (2014).

27. Amad, A. et al. Motor Learning Induces Plasticity in the Resting Brain—Drumming Up a Connection. Cereb. Cortex 27, 2010–2021 (2017).

28. Taubert, M., Lohmann, G., Margulies, D. S., Villringer, A. & Ragert, P. Long-term effects of motor training on resting-state networks and underlying brain structure. NeuroImage 57, 1492–1498 (2011).

29. Sampaio-Baptista, C. et al. Gray matter volume is associated with rate of subsequent skill learning after a long term training intervention. NeuroImage 96, 158–166 (2014).

30. Grefkes, C., Ritzl, A., Zilles, K. & Fink, G. R. Human medial intraparietal cortex subserves visuomotor coordinate transformation. NeuroImage 23, 1494–1506 (2004).

31. Scheperjans, F. et al. Probabilistic Maps, Morphometry, and Variability of Cytoarchitectonic Areas in the Human Superior Parietal Cortex. Cereb. Cortex N. Y. NY 18, 2141–2157 (2008).

32. Culham, J. C. et al. Visually guided grasping produces fMRI activation in dorsal but not ventral stream brain areas. Exp. Brain Res. 153, 180–189 (2003).

33. Goodale, M. A. Action Insight: The Role of the Dorsal Stream in the Perception of Grasping. Neuron 47, 328–329 (2005).

34. Stoodley, C. J., Valera, E. M. & Schmahmann, J. D. Functional topography of the cerebellum for motor and cognitive tasks: an fMRI study. NeuroImage 59, 1560–1570 (2012).

35. Jordan, K., Heinze, H.-J., Lutz, K., Kanowski, M. & Jäncke, L. Cortical Activations during the Mental Rotation of Different Visual Objects. NeuroImage 13, 143–152 (2001).

36. Bushara, K. O. et al. Multiple tactile maps in the human cerebellum: Neuroreport 12, 2483–2486 (2001).

37. Paul, K. I. et al. Mental rotation ability predicts the acquisition of basic endovascular skills. Sci. Rep. 11, 22453 (2021).

38. Bezzola, L., Merillat, S., Gaser, C. & Jancke, L. Training-Induced Neural Plasticity in Golf Novices. J. Neurosci. 31, 12444–12448 (2011).

39. Binkofski, F. & Buccino, G. Motor functions of the Broca’s region. Brain Lang. 89, 362–369 (2004).

40. Wenger, E., Brozzoli, C., Lindenberger, U. & Lövdén, M. Expansion & Renormalization of Human Brain Structure During Skill Acquisition. Trends Cogn. Sci. 21, 930–939 (2017).

41. Trachtenberg, J. T. et al. Long-term in vivo imaging of experience-dependent synaptic plasticity in adult cortex. Nature 420, 788–794 (2002).

42. Xu, T. Rapid formation and selective stabilization of synapses for enduring motor memories. 462, 6 (2009).

43. Braddick, O. J. et al. Brain areas sensitive to coherent visual motion. Perception 30, 61–72 (2001).

44. McMains, S. A. & Somers, D. C. Multiple Spotlights of Attentional Selection in Human Visual Cortex. Neuron 42, 677–686 (2004).

45. Shmuelof, L. & Zohary, E. Dissociation between Ventral and Dorsal fMRI Activation during Object and Action Recognition. Neuron 47, 457–470 (2005).

46. Kipping, J. A. et al. Overlapping and parallel cerebello-cerebral networks contributing to sensorimotor control: An intrinsic functional connectivity study. NeuroImage 83, 837–848 (2013).

47. Ramnani, N. The primate cortico-cerebellar system: anatomy and function. Nat. Rev. Neurosci. 7, 511–522 (2006).

48. Bech, B., Lönn, L., Schroeder, T. V. & Ringsted, C. Fine-motor skills testing and prediction of endovascular performance. Acta Radiol. 54, 1165–1174 (2013).

49. Tardif, C. L. et al. Advanced MRI techniques to improve our understanding of experience-induced neuroplasticity. NeuroImage 131, 55–72 (2016).

50. Lerch, J. P. et al. Studying neuroanatomy using MRI. Nat. Neurosci. 20, 314–326 (2017).

51. Martuzzi, R. et al. A whole-brain voxel based measure of intrinsic connectivity contrast reveals local changes in tissue connectivity with anesthetic without a priori assumptions on thresholds or regions of interest. NeuroImage 58, 1044–1050 (2011).

52. Taubert, M. et al. Dynamic Properties of Human Brain Structure: Learning-Related Changes in Cortical Areas and Associated Fiber Connections. J. Neurosci. 30, 11670–11677 (2010).

53. Kreiser, K. et al. Simulation Training in Neuroangiography—Validation and Effectiveness. Clin. Neuroradiol. 31, 465–473 (2021).

54. Nicholson, W. J. et al. Face and Content Validation of Virtual Reality Simulation for Carotid Angiography: Results from the First 100 Physicians Attending the Emory NeuroAnatomy Carotid Training (ENACT) Program. Simul. Healthc. 1, 147–150 (2006).

55. Saratzis, A., Calderbank, T., Sidloff, D., Bown, M. J. & Davies, R. S. Role of Simulation in Endovascular Aneurysm Repair (EVAR) Training: A Preliminary Study. Eur. J. Vasc. Endovasc. Surg. 53, 193–198 (2017).

56. Oldfield, R. C. The assessment and analysis of handedness: The Edinburgh inventory. Neuropsychologia 9, 97–113 (1971).

57. Taslakian, B., Ingber, R., Aaltonen, E., Horn, J. & Hickey, R. Interventional Radiology Suite: A Primer for Trainees. J. Clin. Med. 8, 1347 (2019).

58. Live Gamer Portable 2 - GC510 | Product | AVerMedia. https://www.avermedia.com/us/product-detail/GC510.

59. R Core Team. R: A Language and Environment for Statistical Computing. (R Foundation for Statistical Computing, 2020).

60. Bates, D., Mächler, M., Bolker, B. & Walker, S. Fitting Linear Mixed-Effects Models using lme4. ArXiv14065823 Stat (2014).

61. Kuznetsova, A., Brockhoff, P. B. & Christensen, R. H. B. **lmerTest** Package: Tests in Linear Mixed Effects Models. J. Stat. Softw. 82, (2017).

62. SPM12 Software - Statistical Parametric Mapping. https://www.fil.ion.ucl.ac.uk/spm/software/spm12/.

63. Ashburner, J. & Friston, K. J. Voxel-Based Morphometry—The Methods. NeuroImage 11, 805–821 (2000).

64. Tournier, J.-D. et al. MRtrix3: A fast, flexible and open software framework for medical image processing and visualisation. NeuroImage 202, 116137 (2019).

65. Andersson, J. L. R. & Sotiropoulos, S. N. An integrated approach to correction for off-resonance effects and subject movement in diffusion MR imaging. NeuroImage 125, 1063–1078 (2016).

66. Jenkinson, M., Beckmann, C. F., Behrens, T. E. J., Woolrich, M. W. & Smith, S. M. FSL. NeuroImage 62, 782–790 (2012).

67. Avants, B. B. et al. A reproducible evaluation of ANTs similarity metric performance in brain image registration. NeuroImage 54, 2033–2044 (2011).

68. Kassambara, A. rstatix: Pipe-Friendly Framework for Basic Statistical Tests. (2021).

69. Neurosynth. https://www.neurosynth.org/.

70. JuBrain Anatomy Toolbox v3.0. (Institute of Neuroscience and Medicine Brain and Behaviour (INM-7), 2021).

