## Supplementary materials for "Visuo-motor transformations in the intraparietal sulcus mediate the acquisition of endovascular medical skill"

**Supplementary Table S1.** The table shows the results from the post-hoc test assessing decreases in intrinsic connectivity in the control group. This post-hoc test followed-up a significant time (pre vs post) * group (experimental vs control group) interaction analysis.

| | | | *Cluster-level* | | | | |
|---|---|---|---|---|---|---|---|
| Decrease in Intrinsic connectivity | SPM Anatomy toolbox | Hemisphere | MNI 152 coordinates | Cluster size | T-value | Z-value | p-value |
| Superior parietal lobule/ aIPS | 24.6% 7A (SPL), 15.7% hlP3 (IPS), 7.7% 5L (SPL), 7.6% 7PC (SPL) | r | 42- 48 60 | 2157 | 8.12 | 6.05 | .000* |
| Superior parietal lobule | 24.6% 7A (SPL), 15.7% hlP3 (IPS), 7.7% 5L (SPL), 7.6% 7PC (SPL) | l | -12 -60 52 | 1128 | 7.11 | 5.56 | .000* |
| Cingulate gyrus / posterior parietal gyrus | 20.6% 5Ci (SPL) | r | 10 -30 40 | 392 | 6.81 | 5.4 | .018 |
| Superior/Middle frontal gyrus / PMC | 25.6% 6d3, 3.9% 6d2 | l | -28 12 54 | 444 | 4.99 | 4.31 | .010* |

Note: r indicates right and l indicates left hemisphere. P-values are shown at cluster-level $p_{FWE}$-corrected < .05. The asterisk * indicates clusters that are still significant when correcting for conducting four post-hoc tests ($p_{FWE}$-corrected < .0125).

**Supplementary Table S2.** The table shows the group (experimental vs control) * time (pre vs post) interaction analysis of the seed-based correlation. The seed is the cluster in the IPS received from the intrinsic connectivity (ICC) analysis.

| | | | *Cluster-level* | | | | |
|---|---|---|---|---|---|---|---|
| Seed-based correlation: IPS with: | SPM Anatomy toolbox | Hemisphere | MNI 152 coordinates | Cluster size | T-value | Z-value | p-value |
| Orbitofrontal cortex | Fo3 | r | 20, 18, -28 | 727 | 6.90 | 5.45 | 0.007* |
| Crus I/ VIIb/VIIIa | FG2 | l | -36, -68, -24 | 933 | 5.93 | 4.90 | 0.002* |
| IPL/aIPS | PFm (IPL) | r | 50, -32, 48 | 1190 | 5.84 | 4.84 | 0.001* |
| IPL/aIPS | PFt/hlP2 | l | -48, -34, 36 | 891 | 5.46 | 4.61 | 0.003* |
| Frontal pole | BA 45 | r | 42, 40, 2 | 701 | 5.33 | 4.53 | 0.008* |
| Cingulate gyrus/precentral gyrus | 5Ci/4a | r | 14, -28, 40 | 759 | 5.04 | 4.34 | 0.006* |
| Inferior temporal gyrus/ Crus I (not in post-hoc test) | FG4 | r | 54, -50, -20 | 490 | 4.87 | 4.23 | 0.031 |

Note: r indicates right and l indicates left hemisphere. P-values are shown at cluster-level $p_{FWE}$-corrected < .05. The asterisk * indicates clusters that are still significant when correcting for conducting four post-hoc tests ($p_{FWE}$-corrected < .0125).

**Supplementary Table S3.** Shows the post-hoc analysis for the contrast experimental group pre- < post-training of the seed-based correlation. The seed is the cluster in the IPS received from the ICC analysis.

| | | | *Cluster-level* | | | | |
|---|---|---|---|---|---|---|---|
| Seed-based correlation: IPS with: | SPM Anatomy toolbox | Hemisphere | MNI 152 coordinates | Cluster size | T-value | Z-value | p-value |
| Orbitofrontal cortex | Fo3 | r | 14, 12, -20 | 533 | 6.76 | 5.37 | 0.024 |
| IPL/aIPS | PFt/hlP2 | l | -46, -40, 36 | 1975 | 5.82 | 4.84 | 0.000* |
| Frontal pole | / | r | 38, 40, 6 | 515 | 6.05 | 4.97 | 0.027 |
| Precentral gyrus | 6mc(SMA)/ 4a | r | 12, -16, 48 | 741 | 5.17 | 4.43 | 0.006* |
| Not in interaction (MTG) | / | l | -60, -40, -4 | 499 | 4.97 | 4.30 | 0.030 |

Note: r indicates right and l indicates left hemisphere. P-values are shown at cluster-level $p_{FWE}$-corrected < .05. The asterisk * indicates clusters that are still significant when correcting for conducting four post-hoc tests ($p_{FWE}$-corrected < .0125).
